# Whole brain optoacoustic tomography reveals strain-specific regional beta-amyloid densities in Alzheimer’s disease amyloidosis models

**DOI:** 10.1101/2020.02.25.964064

**Authors:** Ruiqing Ni, Xose Luis Dean-Ben, Daniel Kirschenbaum, Markus Rudin, Zhenyue Chen, Alessandro Crimi, Fabian F. Voigt, K. Peter R. Nilsson, Fritjof Helmchen, Roger Nitsch, Adriano Aguzzi, Daniel Razansky, Jan Klohs

## Abstract

Deposition of beta-amyloid (Aβ) deposits is one major histopathological hallmark of Alzheimer’s disease (AD). Here, we introduce volumetric multi-spectral optoacoustic tomography (vMSOT), which covers 10×10×10 mm^3^ field-of-view, capable of 3D whole mouse brain imaging. We show for the first time the optoacoustic properties of oxazine-derivative AOI987 probe, which binds to Aβ, and the application of vMSOT for the quantification of brain-wide Aβ deposition. Administration of AOI987 to two common transgenic mouse strains of AD amyloidosis led to a retention of the probe in Aβ-laden brain regions. Co-registered of vMSOT data to a brain atlas revealed strain-specific pattern of AOI987 uptake. A comparison with *ex vivo* light-sheet microscopy in cleared mouse brains showed a good correspondence in Aβ distribution. Lastly, we demonstrate the specificity of the AOI987 probe by immunohistochemistry. vMSOT with AOI987 facilitates preclinical brain region-specific studies of Aβ spread and accumulation, and the monitoring of putative treatments targeting Aβ.

## Introduction

Alzheimer’s disease (AD) is the most common type of dementia ^1, 2^. The abnormal acculumation and spread of amyloid-β (Aβ) deposits play a central role in the pathogenesis of AD and leads to downstream pathophysiological events ^3, 4^. Aβ oligomers, diffuse and fibrillar plaques, can impair neuronal and synaptic function (e.g., long-term potentiation) and selective neuronal loss, and thus contribute collectively to the symptomatology of AD ^5^. Positron emission tomography (PET) imaging of aberrant Aβ deposits with amyloid probes ^11^C-PIB ^6^, ^18^F-flutemetamol ^7^, ^18^F-florbetapir ^8^, ^18^F-fluobetaben ^9, 10^ has been established as diagnostic pathological biomarker for AD pathology in the clinical setting and was included as a new diagnostic criterion ^11^. Higher cortical Aβ loads were reported in the brain from patients with mild cognitive impairment and AD compared to healthy controls using amyloid PET imaging ^6, 8, 9, 10^. PET studies have provided evidence that the accumulation of Aβ deposits are early and initiating events in the pathogenesis of AD ^12^.

Aβ pathology has been successfully modelled in transgenic engineered mouse lines of cerebral amyloidosis, that have proved useful for translational research ^5^. Aβ plaque appearance and spread have been monitored postmortem with histopathology, and resemble in same strains the patterns typically seen in AD patients ^2, 13, 14^. Nevertheless, procedures require extensive lengthy tissue processing and can only be performed cross-sectionally at termination of the experiment*. In vivo* two-photon and optoacoustic microscopic imaging approaches using PIB, methoxy-X04 ^15, 16, 17^, temperature sensitive agents ^18^, Congo red ^19^, and HS-169 ^20^ have enabled monitoring of Aβ growth and disease-modifying therapeutic strategies at the microscopic scale with very high resolution. However, these methods are generally invasive and suffer from a limited field-of-view (FOV) and/or low penetration depth, where Aβ accumulation can only be observed in a small brain region.

Similar to humans, PET imaging of Aβ across the whole brain has been performed in mice using ^11^C-PIB ^21^, ^11^C-AZD2184 ^22^, ^18^F-fluobetaben ^23^, ^18^F-florbetapir ^24^ and labeled antibodies ^25^. While it is a sensitive and quantitative imaging modality, the limited resolution (~1 mm) relative to the mouse brain dimensions (~10×8 mm^2^) hamper to monitor the spread and accumulation of Aβ in smaller brain regions; and the low availability of radiotracer facilities restrict its usage for widespread mouse brain imaging. Optical imaging modalities such as near-infrared fluorescence (NIRF) using amyloid probes AOI987 ^26, 27^, or CRANAD-2 ^28^ and fluorescent molecular tomography (FMT) using AOI987 ^27^ have been shown to provide an alternative for small animal Aβ imaging, but these methods are affected by light scattering and absorption, which limit the achievable spatial resolution and quantification accuracy. Therefore, non-invasive 3D imaging of Aβ deposition across the whole mouse brain with high sensitivity and high spatial resolution has so far not been possible.

Multi-spectral optoacoustic tomography (MSOT) is rapidly evolving to become an established non-invasive imaging modality in preclinical studies involving mice models of many human diseases ^29, 30, 31^. Volumetric MSOT (vMSOT) further enables mapping the bio-distribution of spectrally distinctive contrast agents in entire three-dimensional (3D) regions at multiple spatial and temporal scales ^29, 30, 31^. In contrast to conventional optical imaging, the spatial resolution of vMSOT is not governed by photon scattering but by ultrasound diffraction, thus it can achieve a much higher spatial resolution. As opposed to optoacoustic (OA) microscopy, vMSOT enables high-resolution mapping of endogenous tissue chromophores or exogenous probes at millimeter to centimeter scale depths ^32, 33, 34^. We recently, developed a high-resolution vMSOT systems, which enabled efficient real-time volumetric imaging of the entire mouse brain with spatio-temporal resolution and penetration depth far beyond the limits of optical microscopes ^32, 35^. Herein, we demonstrate the unique capabilities of vMSOT assisted with a NIR oxazine derivative Aβ probe AOI987 ^26^ for *in vivo* whole brain mapping of Aβ deposits in two common mouse models of AD cerebral amyloidosis ^36^.

## Materials and methods

### Animal models

Six transgenic arcAβ mice overexpressing the human APP695 transgene containing the Swedish (K670N/M671L) and Arctic (E693G) mutations under the control of prion protein promoter and five age-matched non-transgenic littermates of both sexes were used in this study (18-24 months-of-age) ^36, 37^. ArcAβ mice are characterized by a pronounced amyloid deposition, cerebral amyloid angiopathy and vascular dysfunction ^14, 38, 39, 40^. Three APP/PS1 mice ^41^ overexpressing the human APP695 transgene containing K670N/M671L and PS1 mutations under the control of prion protein promoter and three age-matched non-transgenic littermates of both sexes (16 months-of-age) were also imaged. Animals were housed in ventilated cages inside a temperature-controlled room and under a 12-hour dark/light cycle. Pelleted food (3437PXL15, CARGILL) and water were provided *ad-libitum*. All experiments were performed in accordance with the Swiss Federal Act on Animal Protection and were approved by the Cantonal Veterinary Office Zurich (permit number: ZH082/18). All procedures fulfilled the ARRIVE guidelines on reporting animal experiments.

#### vMSOT system

The vMSOT set up for *in vivo* mouse brain imaging is illustrated in **Fig. 1a**. It is a further development of functional optoacoustic neuro-tomography (FONT) approach ^32, 35^. Briefly, a short-pulsed (< 10 ns) laser was used to provide an approximately uniform illumination profile on the mouse brain surface with optical fluence < 20 mJ/cm^2^. The excited OA responses were collected with a custom-made spherical array (Imasonic SaS, Voray, France) of 512 ultrasound detection elements with 7 MHz central frequency and > 80 % bandwidth. The array features a central aperture with 8 mm diameter and 3 lateral apertures with 4 mm diameter located at 45° elevation angle and equally spaced (120°) in the azimuthal direction. A custom-made optical fiber bundle (Ceramoptec GmbH, Bonn, Germany) with 4 outputs was used for guiding the laser beam through the apertures of the array. The optoacoustic signals were digitized at 40 megasamples per second with a custom-made data acquisition system (DAQ, Falkenstein Mikrosysteme GmbH, Taufkirchen, Germany) triggered with the Q-switch output of the laser.

**Figure 1.**
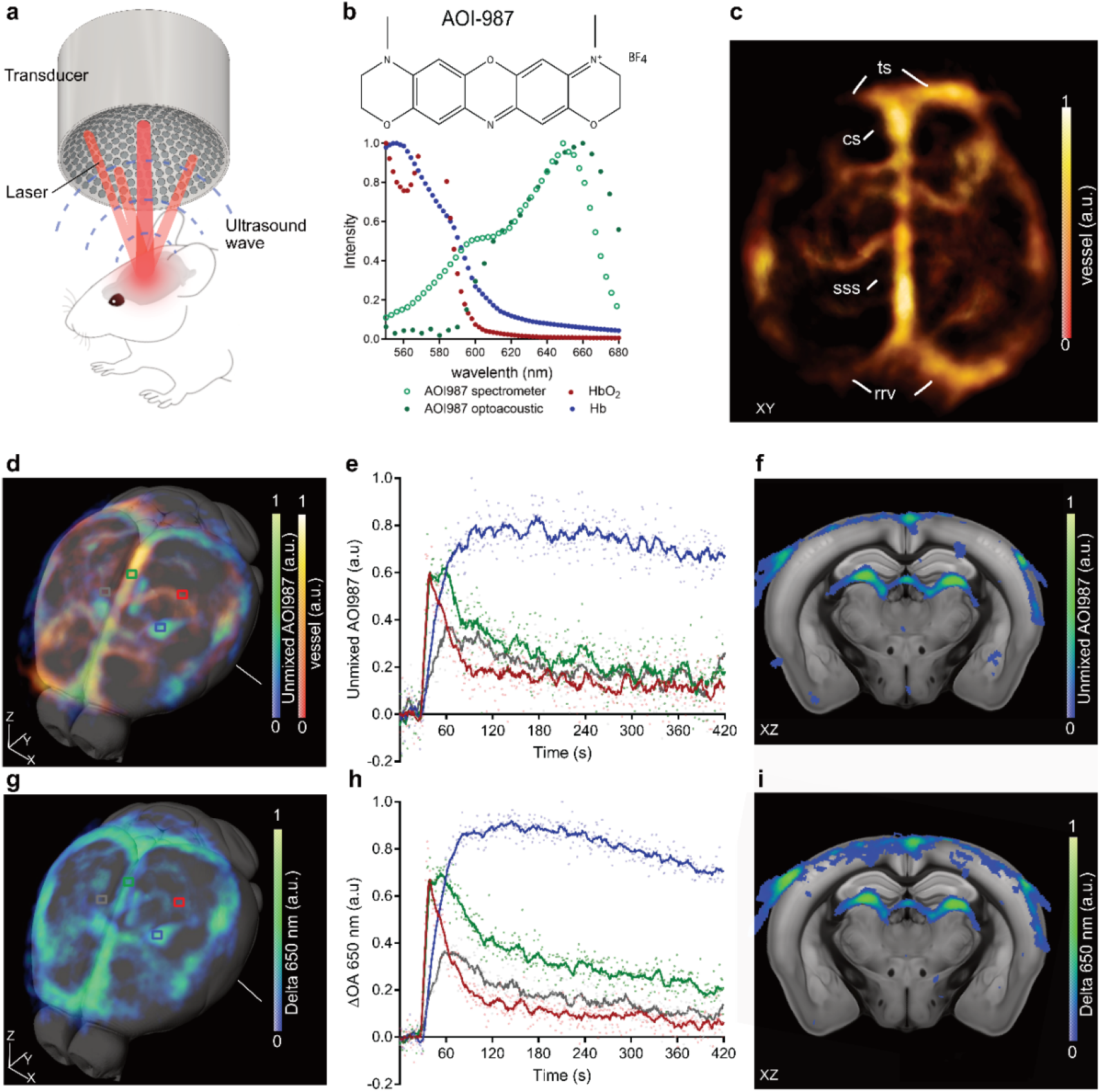
*In vivo* volumetric multispectral optoacoustic tomography (vMSOT) method. **a**) set-up of vMSOT; **b**) absorbance spectrum of AOI987 measured by using vMSOT and using spectrometer; **c**) horizontal view of *in vivo* vMSOT data, vasculature (Hb/HBO_2_, red-yellow); confluence of sinuses (cs), rostral rhinal vein (rrv), superior sagittal sinus (sss), transverse sinus (ts); **d**) 3D rendering of vMSOT data and overlay on MRI structural data with vMSOT signals unmixed for Hb/HbO_2_ corresponding to the vasculature, unmixed for AOI987. AOI987 are indicated in blue-green and Hb/HbO_2_ in red-yellow; **g**) 3D rendering of vMSOT data and overlay on MRI structural data with vMSOT signals for delta absorbance at 650 nm in blue-green; **e**) unmixed AOI987 absorbance; and **h**) delta absorbance at 650 nm from dynamic vMSOT over 0-7 minutes in arcAβ mouse brain. AOI987 injected *i.v*. at 30 seconds. The changes in unmixed AOI987 absorbance and delta absorbance at 650 nm was analyzed in four different brain regions (cortex (red), superior sagittal sinus (blue), hippocampus (green), vessel (grey) on the cortical surface indicated in **d, g**); **f**) unmixed AOI987 absorbance and **h**) delta absorbance at 650 nm in arcAβ mouse brain in one coronal slices and overlay on MRI structural data (anatomical levels are indicated in **d** and **g**).

#### vMSOT of the mouse brains

Mice were anesthetized with an initial dose of 4 % isoflurane (Abbott, Cham, Switzerland) in an oxygen/air mixture (200/800 mL/min), and subsequently maintained at 1.5 % isoflurane in oxygen/air (100/400 mL/min) throughout the measurements. The fur and the scalp over the head of the mice were removed. Animals were placed in prone position on a heating pad that maintains a constant body temperature. Mice were injected intravenously with a 100 μl bolus containing the oxazine probe AOI987 (**Fig. 1b**, 20.5 mg/kg body weight, dissolved in 0.1 M phosphate buffer saline (PBS), pH 7.4) through the tail vein. OA signals were recorded before injection (108 s duration), during injection (432 s duration with *i.v*. injection starting at 30 s after the beginning of acquisition) and 20, 40, 60, 90 and 120 min post-injection (108 s duration each). The pulse repetition frequency (PRF) of the laser was set to 25 Hz and the laser wavelength was tuned between 550 and 680 nm (5 nm step) on a per pulse basis. After the *in vivo* measurements, mice were perfused with 0.1 M PBS (pH 7.4) under ketamine/xylazine/acepromazine maleate anesthesia (75/10/2 mg/kg body weight, i.p. bolus injection) and decapitated. The mouse brains were removed from the skull and imaged *ex vivo* with the vMSOT system. For this, the spherical array was positioned pointing upwards and filled with agar gel to guarantee acoustic coupling, which served as a solid platform to place the excised brains. Uniform illumination of the brain surface was achieved by inserting three arms of the fiber bundle in the lateral apertures of the array and a fourth one providing light delivery from the top. The PRF of the laser was set to 25 Hz and the laser wavelength was tuned between 420 and 680 nm (5 nm step) on a per pulse basis. All recorded OA signals were normalized with the calibrated wavelength-dependent energy of the laser pulse. After the *ex vivo* measurement, mouse brains were fixed in 4 % paraformaldehyde in 0.1 M PBS (pH 7.4) for one day and stored in 0.1 M PBS (pH 7.4) at 4 °C.

#### vMSOT image reconstruction and multi-spectral analysis

vMSOT images were displayed in real time during the acquisition procedure. This was achieved by reconstructing the images with a graphics processing unit (GPU)-based implementation of a back-projection formula ^42^. More accurate image reconstruction was performed off-line. For this, a 3D model-based (iterative) reconstruction algorithm was used, which has been shown to render more quantitative results ^43^. Prior to reconstruction, the collected signals were band-pass filtered with cut-off frequencies 0.1 and 9 MHz. The reconstructed images were further processed to unmix the bio-distribution of AOI987 via per-voxel least square fitting of the OA spectral profiles to a linear combination of the absorption spectra of oxygenated hemoglobin (HbO_2_), deoxygenated hemoglobin (Hb) and AOI987 ^44^. The OA spectra of the probe was experimentally determined as the average spectra of the differential OA image during bolus perfusion at several major vessels in the brain (**Fig. 1b**). The OA spectrum of AOI987 approximately matches the absorption spectrum measured with a spectrophotometer (Avantes BV, Apeldoorn, The Netherlands), being the differences arguably due to the fluorescence of the probe ^45^. The OA spectra of Hb and HbO_2_ was taken from an online database ^46^. The displayed single-wavelength and unmixed images were processed with a median filter with a kernel size 3×3×3.

#### Co-registration with MRI and volume-of-interest analysis

The annotated Allen brain atlas ^47^ was used to identify brain regions in the vMSOT datasets. For this, the reference atlas was manually aligned with the 600 nm vMSOT images, which provided the best contrast for the structure of the cortical vessels of the brain. The same transformation matrix was then applied to the unmixed AOI987 images. Subsequently, these images were co-registered with T_2_-weighted structural MRI images (Ma-Benveniste-Mirrione-T_2_ ^48^) from PMOD (now Bruker). Volume-of-interest (VOI) analysis of 15 brain regions was performed using the embedded Mouse VOI atlas (Ma-Benveniste-Mirrione) in PMOD. Specifically, time course of regional AOI987 absorbance (a.u.) and retention (average of 60-120 min) were calculated. Extra-cranial background signal was removed with a mask from the VOI atlas.

#### Fluorescence microscopy system

A previously established hybrid vMSOT and epi-fluorescence system ^49^ was employed to compare the time course in one arcAβ mouse simultaneously pre- and post *i.v*. bolus injection of AOI987. The laser beam was coupled into the system to provide both the illumination for vMSOT and excitation light for fluorescence imaging. The generated acoustic signal was detected by the ultrasonic matrix array detector and then reconstructed as aforementioned. The excited fluorescence signal was collected by an image guide comprised of 100,000 fibers and then imaged onto an EMCCD camera (Andor Luca R, Oxford Instruments, UK). OA and fluorescence signals were recorded before injection, 20, 40, 60, 90 and 120 min after the injection.

### Brain tissue collection, Staining, confocal imaging and selective plane illumination microscopy

Two arcAβ mice and two non-transgenic littermates were perfused using hydrogel and cleared using a modified electrophoretic clearing chamber for CLARITY ^50, 51^. The detailed procedure for Aβ staining using LCOs (hFTAA and qFTAA) and analysis methods for mesoscopic selective plane illumination microscopy (mesoSPIM) ^52^ are provided as **Suppl materials**. Paraffin fixed brains from arcAβ, APP/PS1 and non-transgenic littermate mice were cut horizontally at 5 μm thickness. Brain sections were stained with anti-Aβ antibody 6E10, AOI987 and fibrillary conformation anti-amyloid antibody OC (details in **Suppl Table 1**) as previously described ^38^. Nuclei were counterstained by 4′,6-diamidino-2-phenylindole (DAPI) while adjacent sections were stained with hematoxylin and eosin (H&E). The slices were imaged at 20× magnification using Pannoramic 250 (3D HISTECH, Hungary) and analyzed using ImageJ (NIH, United States).

The co-localization of the probe AOI987 was evaluated by double staining using antibody 6E10 and OC to amyloid deposits, and confocal microscopy imaging of the cortex, hippocampus and thalamus of arcAβ, APP/PS1 and non-transgenic littermate mice at ×10, ×63 magnification using a Leica SP8 confocal microscope (Leica Microsystems GmbH, Germany) at ScopeM ETH Zurich Hönggerberg core facility. Sequential images were obtained by using 405, 488, 650 nm lines of the laser to excite stained images respectively. Identical settings resolution with Z stack (n = 15) and gain were used. The size of the plaque detected by AOI987 and 6E10 were assessed using full width half maximum (FWHM) method. Double staining using AOI987 and LCOs (hFTAA and qFTAA) were performed to investigate the overlap in detecting Aβ in the mouse brain slices (**Suppl Table 1**).

### Statistics

Group comparison of AOI987 absorbance and immunostaining in multiple brain regions was performed by using two-way ANOVA with Turkey’s *post hoc* analysis (Graphpad Prism, Zurich, Switzerland). All data are presented as mean±standard deviation. Significance was set at \**p* < 0.05.

## Results

### *In vivo* vMSOT amyloid imaging and spectral unmixing

The novel vMSOT system was specifically arranged to cover the entire mouse brain within the FOV of the transducer array (**Fig. 1a)**, approximately 10×10×10 mm^3^. The spatial resolution in a region close to the center of the array is ~ 110 μm. OA signals are generated by endogenous oxygenated (HbO_2_) and deoxygenated (Hb) hemoglobin as well as by the extrinsically administered AOI987 probe. Light absorption in hemoglobin enables distinguishing cerebral vascular structures in the vMSOT images, which serve as an anatomical reference to differentiate different brain regions. The absorption spectra of HbO_2_ and Hb are displayed in **Fig. 1b**. It is shown that hemoglobin absorption strongly decays for wavelengths larger than 600 nm. This decay negatively affects the signal-to-noise ratio of the vMSOT images, but on the other hand enables deeper light penetration. The OA image at 600 nm was found to represent a good trade-off between signal strength and light attenuation for an optimal contrast of the cortical brain vasculature. The maximum intensity projection of the OA image at 600 nm along the depth direction is displayed in **Fig. 1c**, where major vessels and brain areas are labelled (**Suppl video 1**). The OA absorbance spectrum of AOI987 derived from *in vivo* vMSOT imaging data in the brain from arcAβ and APP/PS1 mouse brains is also shown in **Fig. 1b** along with the absorbance measured with a spectrometer (see methods section). The OA spectrum is characterized by a major peak at 650 nm and minor peak at 600 nm. Considering that strong difference in optical attenuation are produced by light absorption in hemoglobin, only wavelengths above 600 nm were considered for spectral un-mixing. A 3D view of the un-mixed image for AOI987 corresponding to 60 min after injection of the probe is shown **SFig. 1** and **Suppl video 2** superimposed to the anatomical image of the brain acquired with MRI (**Fig. 1d**). The same 3D view additionally including the OA image at 600 nm as an anatomical reference is also displayed in **Fig. 1d**.

The accuracy of multispectral un-mixing was assessed by considering the sequence of images acquired during injection of the probe in an arcAβ transgenic mouse. In this case, the bio-distribution of the probe could alternatively be un-mixed by considering the difference with respect to the reference image before injection. For this, the average of 25 OA images corresponding to 650 nm excitation was taken. A comparison of the bio-distribution of AOI987 estimated via difference with respect to the background and via multi-spectral un-mixing is shown in **Figs. 1d, g**. Specifically, one cross-sections of the vMSOT images are displayed superimposed to the corresponding MRI sections (**Figs. 1f, i**). After *i.v*. bolus injection of AOI987 through the mouse tail vein, an OA signal increase was observed in the mouse brain. Immediate uptake was observed in major vessels (superior sagittal sinus and large vessel at the cortical surface), for which the intensity is higher than that in the brain tissue (e.g. in the cortex and hippocampus). The good match between the two un-mixing approaches was further corroborated by considering the time dynamics of the un-mixed signals at specific locations labelled in **Fig. 1d, g**. All the selected profiles showed similar patterns during the 432 s image acquisition window (**Figs. 1e, h**). This validates the results obtained with multi-spectral un-mixing, which is the only available method for other measurements.

### AOI987 optoacoustic signal retention in the brain of arcAβ and APP/PS1 models

The retention of AOI987 was first assessed in the vMSOT and epi-fluorescence images of an arcAβ mouse acquired simultaneously using hybrid system ^49^. Similar temporal profiles the vMSOT and fluorescence signals of the cortical region were observed **(Fig. 1**). Higher retention at 120 min post-injection in the cortex compared to the cerebellum was observed with both imaging systems, suggesting the specific binding of AOI987 in the cortical deposits. The ratio of fluorescence intensity in the cortex/cerebellum at 120 min post-injection was approximately 1.4 in fluorescence microscopy images taken *ex vivo*.

A more comprehensive evaluation of the bio-distribution of the probe was then performed. Three arcAβ mice, three APP/PS1 mice and six non-transgenic littermates were scanned *in vivo* by using vMSOT at several time points until 120 min following AOI987 injection (**Fig. 2a-c**). Multispectral unmixed image of AOI987 channel, and atlas based volume-of-interest analysis were performed (**Fig. 2d**). Two-way ANOVA with Tukey’s *post hoc* analysis showed the interaction between brain region and genotype. Significantly higher AOI987 retention (average of absorbance at 60-120 min post-injection) were observed in the cortex (p = 0.0002), hippocampus (p = 0.0134) and thalamus (p < 0.0001) of arcAβ mice compared to non-transgenic littermates. Only in the cortex (p = 0.0016) of APP/PS1 mice higher AOI987 retention was observed compared to non-transgenic littermates. No apparent accumulation in the brainstem was observed in neither of the transgenic mice. ANNOVA for repeated measure **Fig. 2f-j** showed the time course of AOI987 absorbance in different brain regions of arcAβ, APP/PS1 mice and non-transgenic littermates. The signals were highest in the cortex and thalamus followed by subcortical brain regions and cerebellum and were absent in the brain stem. Fluorescence intensity: F.I.

**Figure 2.**
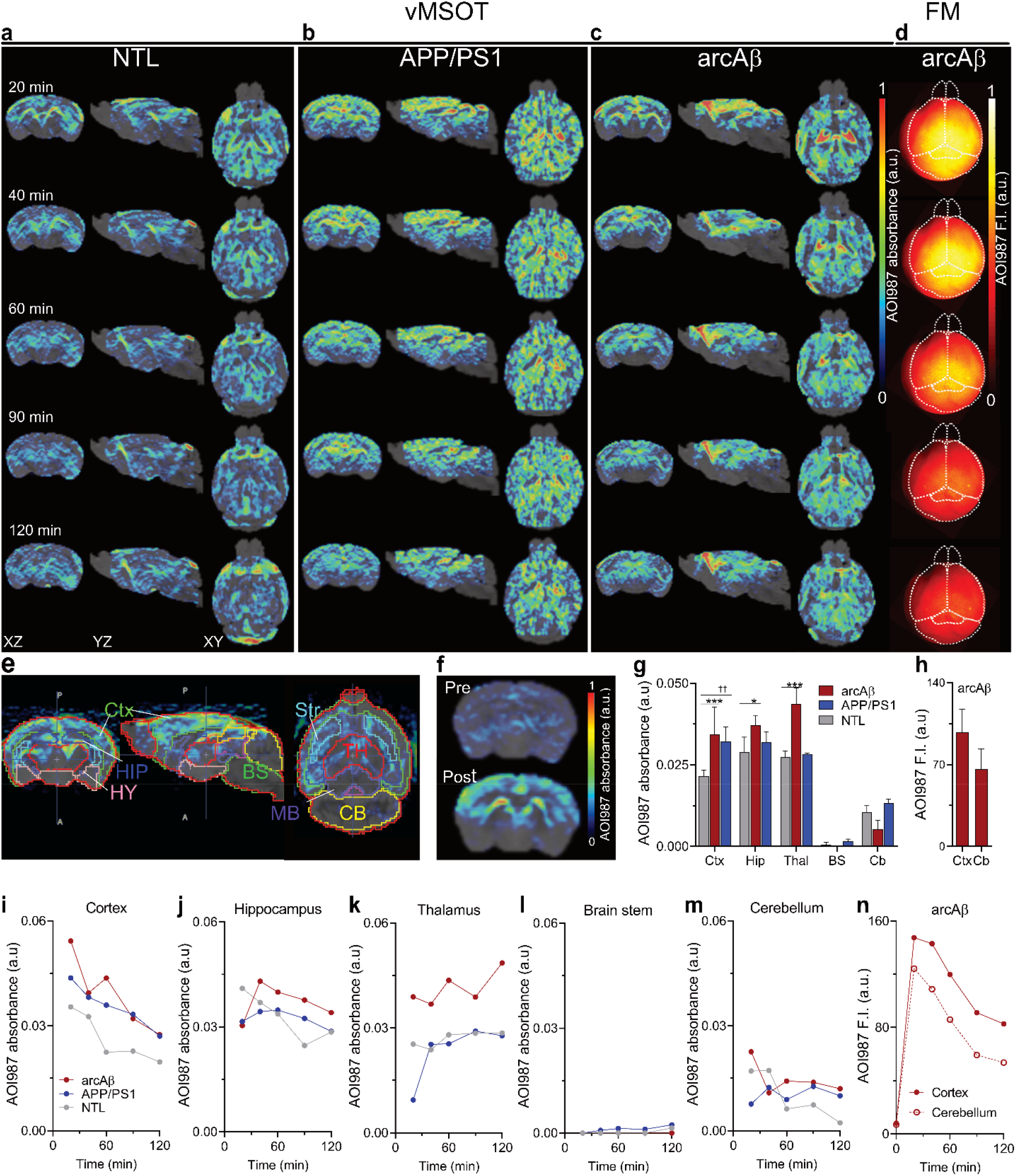
Regional brain AOI987 distribution revealed by using *in vivo* volumetric multispectral optoacoustic tomography (vMSOT) with AOI987 in arcAβ and APP/PS1 mice. vMSOT in **a**) non-transgenic littermates (NTLs), **b**) transgenic APP/PS1 mice and **c**) arcAβ mice; at 20, 40, 60, 90 and 120 min following dye administration showing coronal, sagittal and horizontal views overlaid over the corresponding masked magnetic resonance imaging -based brain atlas. AOI987 absorbance (a.u.) are indicated in rainbow color; **d**) fluorescence microscopy (FM) in one arcAβ mouse at 20, 40, 60, 90 and 120 min following dye administration showing horizontal view; **e**) volume-of-interest used for quantitative analysis of regional AOI987 retention; **f**) pre- and 20 min post dye injection in one arcAβ mouse brain, unmixed, coronal view; **g**) regional comparison of averaged AOI987 absorbance (a.u.) retention at 60-120 min post-injection; **h**) quantification of AOI987 fluorescence intensity in the cortex and cerebellum; **i-m**) time course of cortical, hippocampal, thalamic, brain stem and cerebellar volume-of-interest AOI987 signal (a.u.) in NTL, arcAβ and APP/PS1 mice; **n**) time course of cortical and cerebellar region-of-interest fluorescence intensity (a.u.) in one arcAβ mouse. Brain stem: BS; Cortex: Ctx; Cerebellum: Cb; Hippocampus: Hip; Hy: Hypothalamus; Midbrain MB; Thalamus: TH; \**p* < 0.05; *** *p* < 0.001 comparison between NTL and arcAβ mice, and ^††^*p* < 0.01 comparison between NTL and APP/PS1 mice; two-way ANOVA with *post-hoc* Tukey’s analysis. All data are present as mean ± standard deviation.

### *Ex vivo* vMSOT imaging of the mouse brain

*Ex vivo* vMSOT imaging was performed on PBS perfused (blood washed out) mouse brains excised after *in vivo* imaging. The acquired images served to validate the *in vivo* AOI987 signal bio-distribution in a scenario not affected by cross-talk un-mixing artefacts associated to the presence of Hb and HbO_2_ in a living animal. The same image reconstruction and multi-spectral un-mixing methods used for the analysis of the *in vivo* datasets were considered. The background tissue provide sufficient contrast in the vMSOT for structural information, and hence was used for regional analysis of AOI987 signal in the mouse brain. **Figs. 3a-b** display a comparison of the 3D unmixed AOI987 signal overlaid with structural signal (600 nm) from one non-transgenic mouse and one arcAβ mouse brain. The accumulation of AOI987 signal in the cortex, hippocampus and thalamus of arcAβ mouse suggest specific binding of the probe as the regions are known to express high plaque load ^37^. In contrast, little signals attributed to AOI987 have been observed in the brain of a non-transgenic littermate (**Fig. 3c, SFig. 2**).

**Figure 3.**
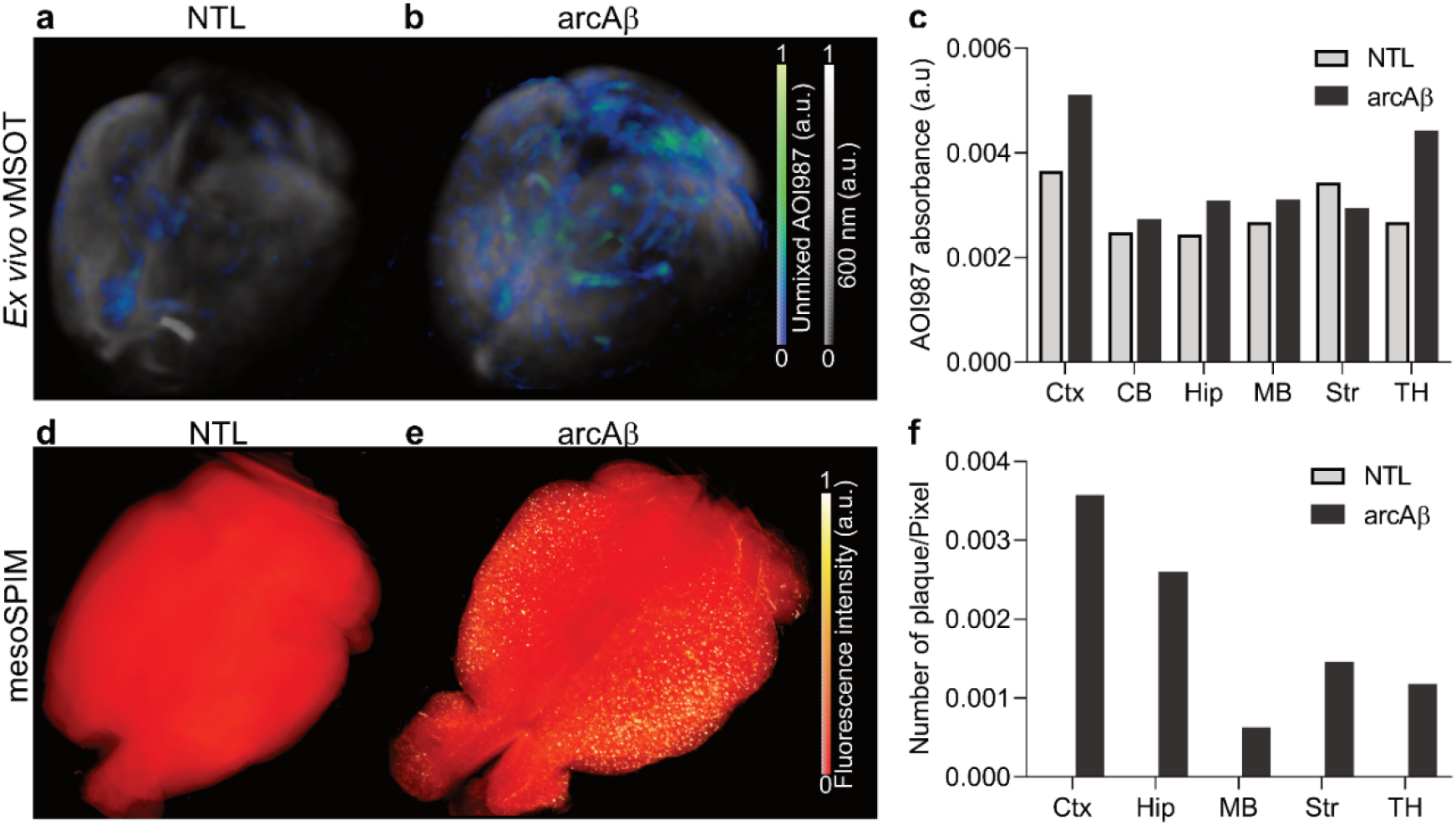
*Ex vivo* volumetric multispectral optoacoustic tomography (vMSOT) imaging and whole brain light-sheet microscopy using mesoSPIM of brain from transgenic arcAβ and non-transgenic littermate (NTLs) mice. **a)** 3D rendering of vMSOT data overlay of vMSOT structural information with vMSOT signals unmixed for AOI987 in one non-negative control mouse; and **b**) one arcAβ mouse; AOI987 are indicated in blue-green; 600 nm vMSOT signal provide structural information (a.u.) of the brain at *ex vivo*; **c**) Quantification of *ex vivo* regional AOI987 absorbance (a.u.) in NTL and arcAβ mouse brain; **d**) 3D rendering of mesoSPIM data of amyloid-beta distribution (stained by using luminescent conjugated oligothiophene) in one NTL mouse; and **e**) one arcAβ mouse. **f**) Quantification of regional fluorescence intensity in NTL and arcAβ mouse brain. Cortex: Ctx; Hippocampus: Hip; Midbrain: MB; Striatum: Str; Thalamus: TH.

### MesoSPIM for mapping amyloid distribution in the whole brain of arcAβ mouse

MesoSPIM ^52^ has been used to determine the 3D distribution of AOI987 in brains of arcAβ mice and non-transgenic littermates with luminescent conjugated oligothiophene probe. SPIM imaging of the cleared brain of arcAβ mice revealed signals in the cortical, hippocampal and thalamic parenchyma as well as in vascular structures, indicative of aggregated Aβ accumulation. In contrast, brains of non-transgenic littermates displayed only background signal intensity with the exception of few high intensity signal attributed to vessels **(Figs. 3d-e**, **SFig. 3 and Suppl video. 3**). Regionally resolved quantitative analysis of numbers of amyloid deposits revealed a higher load in the cortical, hippocampus and thalamic regions-of-interest as compared to cerebellar and other subcortical regions (**Fig. 3f**, **Suppl materials**). Regions were defined on the basis of the Allen brain atlas. Double staining using AOI987 and hFTAA shows predominantly overlap in the detection of both vascular and parenchymal plaque in the arcAβ mouse brain (**SFigs. 4e-k**), whereas less convergence between AOI987 and qFTAA (**SFigs. 4a-d)**.

### Immunohistochemical staining and confocal microscopy

Immunohistochemical staining were performed on horizontal brain tissue sections from arcAβ, APP/PS1 mice and non-transgenic littermates using AOI987, 6E10, OC and counterstained using DAPI (**Fig. 4**). In APP/PS1 mice, the Aβ plaques were predominantly observed in the parenchyma, most pronounced in the cortex (**Figs. 4i-j)** and hippocampus (**Figs. 4e-f**). In arcAβ mice, the 6E10 and AOI987 stained Aβ deposits were distributed both in the cortical (**Figs. 4k-l**), hippocampal (**Figs. 4g-h**) and thalamic (**Figs. 4o-p**) parenchyma as well as along vascular structures predominantly in the cortex. The staining distribution reveals that AOI987 and 6E10 interact with both parenchymal and vascular amyloid deposits, the latter being indicative of cerebral amyloid angiopathy. The plaques appeared less diffuse in arcAβ mice compared to that in APP/PS1 mice. The size of deposits in the cerebral cortex stained by AOI987 tends to be smaller than that stained by 6E10, the difference being more pronounced in APP/PS1 than in the arcAβ mice (**Fig. 4u**). For smaller plaques, AOI987 and 6E10 almost fully overlapped (**Fig. 4q**). Similarly, there was a good correspondence of OC stained fibrillar Aβ with AOI987 in the cortical regions-of-interest of APP/PS1 mice (**Fig. 4r**) and arcAβ mice (**Figs. 4s-t).** No signal of AOI987 was detected in the brain sections from non-transgenic littermates as there is no amyloid plaque (**Figs. 4v-w**).

**Figure 4.**
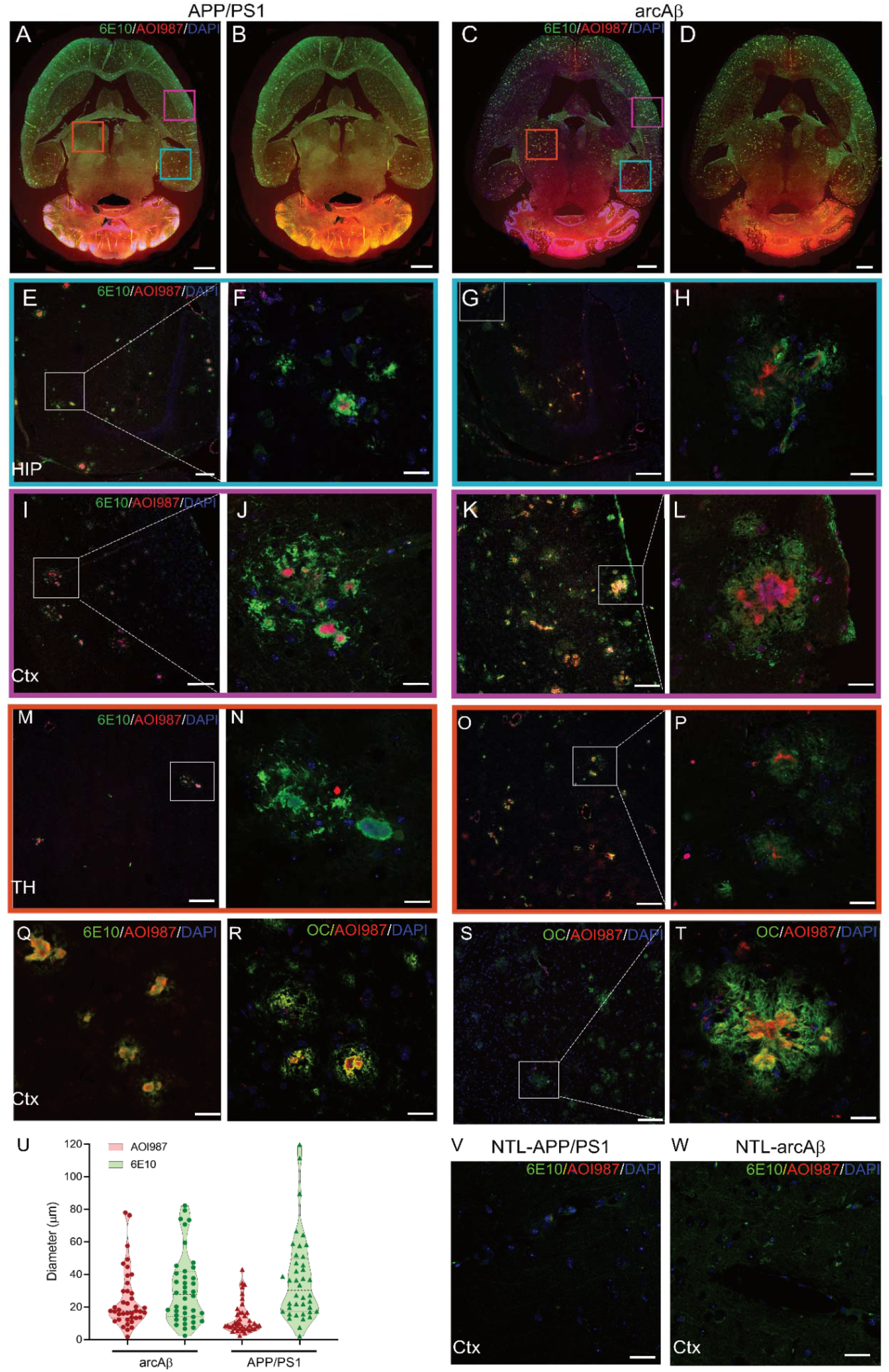
Staining for Aβ deposition in APP/PS1 and arcAβ mouse brain tissue sections. (**a, b**) confocal microscopic images of horizontal whole brain sections from APP/PS1 mouse; and (**c, d)** from arcAβ mouse; DAPI (blue), Alexa488-6E10 (green), AOI987 (red); Zoomed-in images are shown for hippocampus (**e-h**, blue square in (**a, c**)); cortex (**i-l, q**, **s** magenta square in (**a, c**)) and thalamus (**m-p**, orange square in (**a, c**)), respectively, demonstrating the co-localization of 6E10 and AOI987 to amyloid-beta plaque in APP/PS1 and arcAβ mouse brain with further magnification of selected regions-of-interest (white squares). DAPI (blue), Alexa488-6E10 (green), AOI987 (red); **r-t**) Confocal microscopic images of cortical section from APP/PS1 and arcAβ mice indicates co-localization of OC targeting amyloid fibrils and AOI987 stained deposits, OC (green), AOI987 (red), DAPI (blue); Again, volume of AOI987 stained structures appears smaller than OC stained ones; (**u**) size analysis indicates that stained volume of cortical parenchymal plaques was larger when using 6E10 (targeting Aβ_1-16_) as compared to AOI987 stains for both APP/PS1 and arcAβ mice; (**v, w**) confocal imaging in horizontal cut cortex from non-transgenic littermate mice, DAPI (blue), Alexa488-6E10 (green), AOI987 (red). Scale bar =1 mm (**a-d**); 100 μm (**e, g, i, k, m, o, s**); 20 μm (**f, h, j, l, n, p, r, t, v, w**) respectively.

## Discussion

The development of new tools for non-invasive detection of Aβ deposits with high resolution in animal models of AD is critical for a better understanding of the disease mechanisms and translational development of Aβ-targeted therapies. We developed a vMSOT system used previously for FONT^32, 35^, for fast 3D whole brain amyloid imaging. AO1987, an Aβ-targeting probe ^26, 27^, for optoacoustic imaging in two different models of AD amyloidosis. Fast volumetric acquisition of multispectral optoacoustic tomographic data together with a novel spectral unmixing approach and brain atlas-based data co-registration allowed us to systematically quantify brain-wide distribution of Aβ plaque pathology at mesoscale resolution. The comparison of two commonly used models of AD amyloidosis with vMSOT enabled to reveal strain-specific pattern of Aβ deposition.

So far, at high-resolution imaging of Aβ load across the whole brain is only possible *ex vivo* by histopathological and microscopic techniques ^13, 14^ that require extensive tissue processing and are both painstaking and time-consuming. The vMSOT employed in this work is capable of imaging the entire mouse brain at mesoscale resolution (~110 μm) and is sensitive to optical contrast agents targeting Aβ. Herein, we demonstrated that 3D vMSOT imaging assisted with the amyloid-binding probe AOI987 accurately maps Aβ deposits across the murine brain in two transgenic mouse models of cerebral amyloidosis commonly used in AD research. In arcAβ mice intracellular punctate Aβ deposits start concomitantly with robust cognitive impairments at the age of 6 months before the onset of Aβ plaque formation and cerebral amyloid angiopathy ^36^. In APP/PS1 mice, the Aβ deposits accumulate earlier and mainly in parenchyma ^41^. We found that vMSOT mapping of AOI987 specifically detected high cortical, hippocampal and thalamic retention in arcAβ mice, while only higher cortical retention in APP/PS1 mice. The *in vivo* Aβ distribution patterns revealed with vMSOT were shown to be in accordance with our immunohistochemical staining results and the known Aβ distribution in both transgenic mouse models ^36, 41^.

The limitations of *ex vivo* Aβ assessment techniques has spurred the search for Aβ imaging techniques. PET would be the method of choice, as it would directly facilitate translational studies assessing Aβ in AD patient studies ^6, 7, 8, 10^ ^9, 10^ and animal models ^21^ ^22^ ^23^. In transgenic models PET amyloid imaging is limited as currently achievable microPET for small animal achieves ~1 mm resolution. vMSOT provides ~100 μm spatial resolution which is 10× higher. This becomes significant when Aβ load is assessed in different brain regions. While PET can only coarsely discriminate between cortical and subcortical structures, novel vMSOT can assess Aβ accumulation in different regions. This advantage helps to identify mouse strain-specific pattern of Aβ deposition and to study the spread of Aβ across the brain over the disease course.

vMSOT further covers a larger range of spatial scales than pure optical methods, which either lack the penetration depth (i.e. optical microscopy) or the spatial resolution (i.e. diffuse optical imaging and tomography) ^29^. AO1987 has been used in NIRF imaging and FMT ^26, 27^ for detecting amyloid pathologies in an APP23 mouse model of AD cerebral amyloidosis. In NIRF imaging, the fluorescence signal that is quantified over the brain regions contains also other signal contribution from superficial tissues, i.e. skin, muscle and skull, rending quantification difficult. Indeed, biological tissues are highly scattering and light propagation is mainly diffusive for depths beyond 1 mm. Although mathematical models based on diffuse optics are available e.g. for FMT ^27^, quantification is generally not possible in the diffuse regime of light. In contrast, due to the true 3D mapping capability of vMSOT, the photoacoustic signal can be quantified for the different compartments provided proper image reconstruction and processing is performed. Thus, the results obtained indicate that the unprecedented capabilities provided empower vMSOT as a new method that can clearly outperform existing *in vivo* amyloid imaging technologies in animal models.

In the current implementation, a linear un-mixing algorithm was used for calculating the bio-distribution of AOI987 in deep brain regions such as the hippocampus and the thalamus. However, it is noted that the wavelength-dependent optical attenuation causes distortion in the OA spectra of signals generated at deep locations (spectral colouring), while the non-uniform laser beam shape further affects the superficial signals. These and other effects lead to cross-talk artefacts in the un-mixed image corresponding to the contrast agent of interest, so that more advanced algorithms are required for an optimal performance. The good correspondence between the *in vivo* and *ex vivo* regional bio-distribution however suggests that this un-mixing approach is generally suitable for our purposes. The reconstruction algorithm is also essential for quantifying the concentration of the probe. The model-based algorithm employed to reconstruct the images off-line has been shown to provide more quantitative results than standard back-projection algorithms. However, acoustic distortions associated to speed of sound heterogeneities, acoustic scattering and attenuation as well as the response of the ultrasound sensing elements of the array are known to additionally play a role in the quality of the images rendered. More accurate estimation of the bio-distribution of the contrast agent of interest can also be facilitated with pharmacokinetic modeling, which has been previously developed based on the anatomical information in cross-sectional MSOT images ^53, 54^. Registration between MSOT and MRI/atlas can additionally boost the performance of these models and further provide a better anatomical reference for regional analysis. Automatic registration methods have been established for 2D dataset ^55, 56^, which can potentially be adapted to 3D vMSOT datasets. However, as both arcAβ and APP/PS1 mice showed amyloid deposits in the cerebellum at an old age, regions typically considered as reference region, we therefore opted for average absorbance retention (60-120 min) rather than the reference tissue region modelling. Alternatively, brainstem regions might be considered as reference as they hardly display probe accumulation.

Our approach aimed to provide a consist plaque labeling method using a single Aβ-targeted probe and standardized imaging conditions to quantitatively compare plaque density across individual animals and strains. However, we noted that the size of deposits in the cerebral cortex stained by AOI987 was slightly smaller than that stained by 6E10, the difference being more pronounced in APP/PS1 than in the arcAβ mice. We are still confident that vMSOT does not grossly underestimate plaque load, which is similar to what has been reported with histopathology and microscopy ^16, 57^.

Aβ deposition in the brain is complex process and involves various molecular species including Aβ oligomers, Aβ fibrils, as well as large insoluble aggregates (plaques). Aβ peptides are generated via sequential proteolytic cleavage of the amyloid precursor protein (APP) ^5^. Initial cleavage of APP by β-secretase generates a membrane-bound C-terminal fragment of APP (β-CTF), which is sequentially processed by γ-secretase resulting in the formation of longer species, like Aβ40 and Aβ42/43, in addition to several other C-terminally truncated Aβ variants ^58^. A full set of Aβ isoforms with both N- and C-terminally truncated Aβ species can be detected in the post-mortem AD brain and cerebrospinal fluid ^59^. N-terminally truncated Aβ isoforms show an increased aggregation propensity and have previously been associated with the deposition of dense-core amyloid plaques in the brain ^60^. In contrast, C-terminally truncated shorter Aβ isoforms show a higher solubility *in vitro* and are more commonly detected in the cerebral vasculature ^61^. The high degree of heterogeneity in the molecular architecture of misfolded Aβ observed in patients with AD may be related to the diversity of disease-associated mutations ^62^. LCO has been applied to detect Aβ β-sheet structures in different animal models of AD amyloidosis ^63, 64^ and recognizing structural variants of Aβ fibrils from brain tissue from AD patients carrying different mutations ^62, 65^. Yet the approach is not suited for in *vivo* applications. Instead, AOI987 has been used due to its strong absorption in the far-red/NIR range warranting efficient tissue penetration of the excitation light, its water solubility and its high binding affinity to Aβ fibrils ^26^. Moreover, the dye has been shown to efficiently cross the blood-brain barrier passing property enabling NIRF imaging and FMTapplications ^26^. We observed regional plaque density with vMSOT in our study for arcAβ and APP/PS1 mice. An earlier study compared five mouse lines displaying cerebral amyloidosis, i.e. arcAβ, APP23, APPswe, PSAPP and ΔE22 mice for two different PET tracers ^11^C-PIB and ^18^F-flutemetamol ^66^. The differences across mouse lines may arise due to regional differences in Aβ deposition, due to structural variations of the Aβ species with binding sites exhibiting different affinity to the respective imaging probe, as well as different binding sites for different imaging probes or even several binding sites per Aβ molecular entity ^62, 63, 65, 67, 68^. The current *in vivo* data do not allow discriminating these cases. Further studies will be needed to elucidate if AOI987, PIB and flutemetamol behave differently, i.e. reveal slightly different results regarding the spatial distribution, and different Aβ deposits.

3D amyloid imaging with light sheet microscopy in cleared brain tissues (3DISCO or CLARITY^51^) of AD mouse models using anti-Aβ antibodies, Congo-red and methoxy-X04 ^69^ has been reported previously. While the method enables imaging at single-cell resolution in the entire brain ^50^ it requires extensive tissue processing and lengthy image post-processing limiting the capability of high through-put imaging. Herein, we have used *ex vivo* light-sheet microscopy to assess the distribution of Aβ plaques in the arcAβ mouse brain, which confirmed our findings from vMSOT mapping of parenchymal and vascular Aβ deposits. Hence, vMSOT in combination with an Aβ specific absorbing dye such as AOI987, enables fast volumetric mapping of Aβ distribution across the whole mouse brain *in vivo* combining the high sensitivity of optical imaging with the high spatial resolution of ultrasound imaging.

In conclusion, we demonstrated the feasibility of quantitative whole brain imaging of the Aβ distribution in mouse models of AD amyloidosis at high spatial and temporal resolution *in vivo* by vMSOT. Brain MRI atlas was used as a structural reference. *In vivo* results have been validated using light-sheet microscopy of cleared brain specimen and immunohistology. The vMSOT imaging platform lends itself for longitudinal monitoring of the disease process, its modulation by therapeutic interventions targeting at Aβ, and potentially for studying other proteinopathy (such as tauopathy) in animal models ^70^.

## Availability of data and material

The datasets generated and/or analyzed during the current study are available in the repository zenodo DOI 10.5281/zenodo.3576174. The light-sheet microscopy dataset and github code for registration and quantification with Allen brain atlas are available at github (details in **Suppl materials**).

## Author Contributions

RN, LD, DR, JK conceived and designed the study; RN, LD, DK performed the experiments; MR provided AOI987; KPRN provided FTAAs. RN, LD, AC, DK analyzed the data; RN, LD, DR and JK interpreted the results; RN, LD, MR, DR, and JK wrote the paper; all coauthors contributed constructively to the manuscript.

## Acknowledgement

The authors acknowledge technical support from Dr. Linjing Mu at Institute of Pharmaceutical Sciences, ETH Zurich, Dr. Joachim Hohl, Dr. Gabriella Bodizs of the Scientific Center for Optical and Electron Microscopy (ScopeM) of ETH Zurich; Ms Gloria Shi, Ms Marie Rouault at Institute for Biomedical Engineering of ETH Zurich; Dr. Luka Kulic, Ms Priyanka Ravikumar at Institute for Regenerative Medicine of University of Zurich for technical help.

